# A YTHDF-PABP axis is required for m^6^A-mediated organogenesis in plants

**DOI:** 10.1101/2023.07.03.547513

**Authors:** Mathias Due Tankmar, Marlene Reichel, Laura Arribas-Hernández, Peter Brodersen

## Abstract

*N6*-methyladenosine (m^6^A) in mRNA is key to eukaryotic gene regulation. Many m^6^A functions involve specialized RNA-binding proteins that recognize m^6^A via a YT521-B Homology (YTH) domain. YTH domain proteins contain long intrinsically disordered regions (IDRs) that may mediate phase separation and interaction with protein partners, but whose precise biochemical functions remain largely unknown. The *Arabidopsis thaliana* YTH domain proteins ECT2, ECT3 and ECT4 accelerate organogenesis through stimulation of cell division in organ primordia. Here, we focus on ECT2 to reveal molecular underpinnings of this function of ECT2/3/4. We show that stimulation of leaf formation requires the long N-terminal IDR, and we identify two short IDR-elements required for ECT2-mediated organogenesis. Of these two, a tyrosine-rich 19-amino acid region is necessary for binding to a small subset of proteins that includes the major cytoplasmic poly(A)-binding proteins PAB2, PAB4 and PAB8. Remarkably, overexpression of PAB4 in leaf primordia partially rescues the delayed leaf formation in *ect2 ect3 ect4* mutants, suggesting that the ECT2-PAB2/4/8 interaction on target mRNAs of organogenesis-related genes may overcome limiting PAB concentrations in primordial cells.

## INTRODUCTION

Methylation of internal adenosine in pre-mRNA to form *N6*-methyladenosine (m^6^A) is a key regulatory step in eukaryotic mRNA biogenesis and functionality. In both plants and animals, it may dictate distinct processing steps in the nucleus (Haussmann *et al*, 2016; Lence *et al*, 2016; Parker *et al*, 2020; Pontier, 2019; Roundtree *et al*, 2017; Tang *et al*, 2018; Wei *et al*, 2021; Zheng *et al*, 2013), and influence properties such as mRNA half-life and translatability in the cytoplasm (Herzog *et al*, 2017; Ke *et al*, 2017; Meyer *et al*, 2015; Sommer *et al*, 1978; Wang *et al*, 2014; Wang *et al*, 2015; Weng *et al*, 2018; Zaccara & Jaffrey, 2020). The main readers of m^6^A contain a YT521-B homology (YTH) domain (Dominissini *et al*, 2012; Patil *et al*, 2018; Wang *et al*., 2014; Wang *et al*., 2015) that recognizes m^6^A through a highly conserved aromatic cage (Li *et al*, 2014b; Luo & Tong, 2014; Theler *et al*, 2014; Xu *et al*, 2015; Xu *et al*, 2014; Zhu *et al*, 2014). YTH domains fall into two main phylogenetic groups, DC and DF (Patil *et al*., 2018). YTHDC proteins are often nuclear (Nayler *et al*, 2000; Zhang *et al*, 2010) and typically contain long intrinsically disordered regions (IDRs) in addition to the YTHDC domain (Patil *et al*., 2018). In some cases, YTHDC domains are part of multi-domain proteins, such as the long isoform of the plant Cleavage and Polyadenylation Specificity Factor 30 (CPSF30) implicated in selection of poly(A) sites (Hou *et al*, 2021; Pontier, 2019; Song *et al*, 2021). YTHDF proteins are typically cytoplasmic (Arribas-Hernández *et al*, 2018; Arribas-Hernández *et al*, 2021b; Kan *et al*, 2021; Wang *et al*., 2014; Worpenberg *et al*, 2021; Zhang *et al*., 2010) and have a YTH domain placed at the C-terminus after an IDR of variable length (Meyer, 2017). Plant YTHDF proteins are, for this reason, referred to as EVOLUTIONARILY CONSERVED C-TERMINAL REGION (ECT) (Ok *et al*, 2005), and higher plants encode an expanded family of YTHDF proteins with 11 in *Arabidopsis thaliana* (arabidopsis) compared to 3 in mammals, and 1 in *Drosophila* and yeast (Fray & Simpson, 2015; Li *et al*, 2014a).

In arabidopsis, the YTHDF proteins ECT2, ECT3 and ECT4 are specifically expressed in rapidly dividing primed stem cells in organ primordia (Arribas-Hernández *et al*., 2018; Arribas-Hernandez *et al*, 2020). ECT2-4 are necessary for the rapid division of these cells, such that organogenesis is delayed due to slow growth in *ect2-1 ect3-1 ect4-2* (henceforth abbreviated *te234*) mutants (Arribas-Hernández *et al*., 2018; Arribas-Hernandez *et al*., 2020). Similar developmental defects are observed in plants with hypomorphic mutations in m^6^A methyltransferase components or in which such components are knocked down during post-embryonic development (Bodi *et al*, 2012; Růžička *et al*, 2017; Shen *et al*, 2016). ECT2 and ECT3 are the main m^6^A effectors in rapidly dividing primordial cells, since *ect2 ect3*, but no other double mutant examined, exhibits clear delays in organogenesis (Arribas-Hernández *et al*., 2018; Flores-Téllez *et al*, 2023; Martínez-Pérez *et al*, 2022). Accordingly, the involvement of ECT4 in organogenesis is only revealed by exacerbation of the organogenesis defects in *ect2 ect3* mutants upon additional inactivation of *ECT4* (Arribas-Hernández *et al*., 2018; Arribas-Hernandez *et al*., 2020).

ECT2 binds directly to m^6^A deposited in DRACH or GGAU contexts in target mRNAs by the nuclear MTA/MTB adenosine methyl transferase complex (Arribas-Hernández *et al*, 2021a; Parker *et al*., 2020), and all described *in vivo* functions of ECT2 and ECT3 depend fully on intact m^6^A-binding pockets (Arribas-Hernández *et al*., 2018; Arribas-Hernandez *et al*., 2020; Martínez-Pérez *et al*., 2022). The genetic redundancy between ECT2 and ECT3 is reflected in their strongly overlapping sets of targets, for which they compete to bind *in vivo* (Arribas-Hernández *et al*., 2021b). In the absence of ECT2, ECT3 and ECT4, target mRNAs of ECT2/3 tend to be downregulated (Arribas-Hernández *et al*., 2021b), perhaps because of more rapid mRNA degradation, although indirect effects acting at the level of target mRNA transcription cannot be excluded at this point. Consistent with a stabilizing effect of ECT2/3 binding to m^6^A, increased mRNA degradation rates upon global inhibition of transcription have been observed for a few m^6^A targets in plants with post-embryonic methyltransferase knockdown (Anderson *et al*, 2018). Because sequence signatures consistent with endonucleolytic cleavage were detected in these transcripts (Anderson *et al*., 2018), it is possible that ECT2/3 binding to m^6^A in target mRNA exerts merely a passive, protective role via m^6^A-binding mediated by the YTH-domain. At the extreme, such a model would be compatible with ECT proteins relying nearly exclusively on their YTH domains for *in vivo* function, with the IDRs perhaps mainly there to drive sequestration into distinct biophysical phases in response to stress, as shown previously to be the case (Arribas-Hernández *et al*., 2018; Scutenaire *et al*, 2018). Several lines of evidence suggest, however, that the IDR plays a more active and specific role in the mechanism underlying m^6^A function. First, individual nucleotide resolution crosslinking-immunoprecipitation (iCLIP) experiments with ECT2 recovered sequence tags attributable to the IDR, suggesting a direct involvement of the IDR in RNA binding (Arribas-Hernández *et al*., 2021a). Second, systematic tests of the ability of arabidopsis ECT paralogs to substitute for ECT2-4 upon ectopic expression in leaf primordia identified three paralogs (ECT1, ECT9 and ECT11) with divergent function, and showed that the IDRs of these proteins are major contributors to the functional divergence (Flores-Téllez *et al*., 2023). It is an obvious possibility that the IDR mediates contacts to proteins relevant for mRNA translation and stability. Precedents for such a model come from studies of human YTHDF2 in which different regions of the N-terminal IDR interact with (i) the CNOT1 subunit of the CCR4-NOT complex (Du *et al*, 2016), (ii) the adaptor protein HRSP12, which acts as a bridge between YTHDF2 and the RNAse P/MRP complex (Park *et al*, 2019), and (iii) the nonsense-mediated mRNA decay factor UPF1 (Boo *et al*, 2022). Thus, if a similar conceptual model for YTHDF function applies to plant ECT proteins, distinct IDR elements crucial for *in vivo* function may be defined. Here, we use ECT2 to address this basic question regarding YTHDF function in plants. We show that the first 410 amino acid residues (aa) in the N-terminal IDR are indispensable for function *in vivo* and, surprisingly, that this region is as important for RNA association *in vivo* as the m^6^A-binding pocket in the YTH domain. We also identify two distinct, ∼20-40 aa elements that are required for the molecular functions of ECT2 in plant development. One of these elements has little, if any effect on RNA association, but is required for association with the major cytoplasmic poly(A) binding proteins PAB2/4/8 *in vivo*. The biological importance of the physical ECT2-PAB association is supported by the suppression of leaf formation defects in *te234* mutants upon overexpression of PAB4 in leaf primordia. These observations support a model in which ECT2/3-PAB2/4/8 interaction on target mRNAs of organogenesis-related genes overcomes limiting concentrations of PABP in rapidly dividing primordial cells.

## RESULTS

### Experimental system for assessment of ECT2 function in vivo

The *te234* mutant displays a consistent 2-day delay in emergence of the first true leaves compared to wild type (Arribas-Hernández *et al*., 2018). This phenotype is easily scored at 9 days after germination (DAG) in which wild type has clearly visible, > 1 mm long true leaves while the first true leaves of *te234* are barely visible and always < 0.5 mm in size (Fig 1A, left panel). This clear and fully penetrant phenotype is restored by the transgenic expression of *ECT2-mCherry* driven by the endogenous *ECT2* promoter (Arribas-Hernández *et al*., 2018), or by the *RPS5A/US7Y* (At3g11940) promoter (Flores-Téllez *et al*., 2023) with activity in highly dividing meristematic cells (Scarpin *et al*, 2023; Weijers *et al*, 2001). Since *te234* mutants are fertile and can be transformed with normal efficiency, the ability to complement the delayed leaf formation of *te234* at 9 DAG constitutes a sensitive readout for *ECT2* function *in vivo*. We previously used this system to show that eight of the eleven arabidopsis YTHDF proteins have molecular functions sufficient to complement the leaf formation defects of *te234* mutants when expressed in organ primordia (Flores-Téllez *et al*., 2023). Thus, we set out to use this robust system to test a series of IDR-deletion mutants in ECT2 to identify elements of functional importance (Fig 1A).

**Figure 1.**
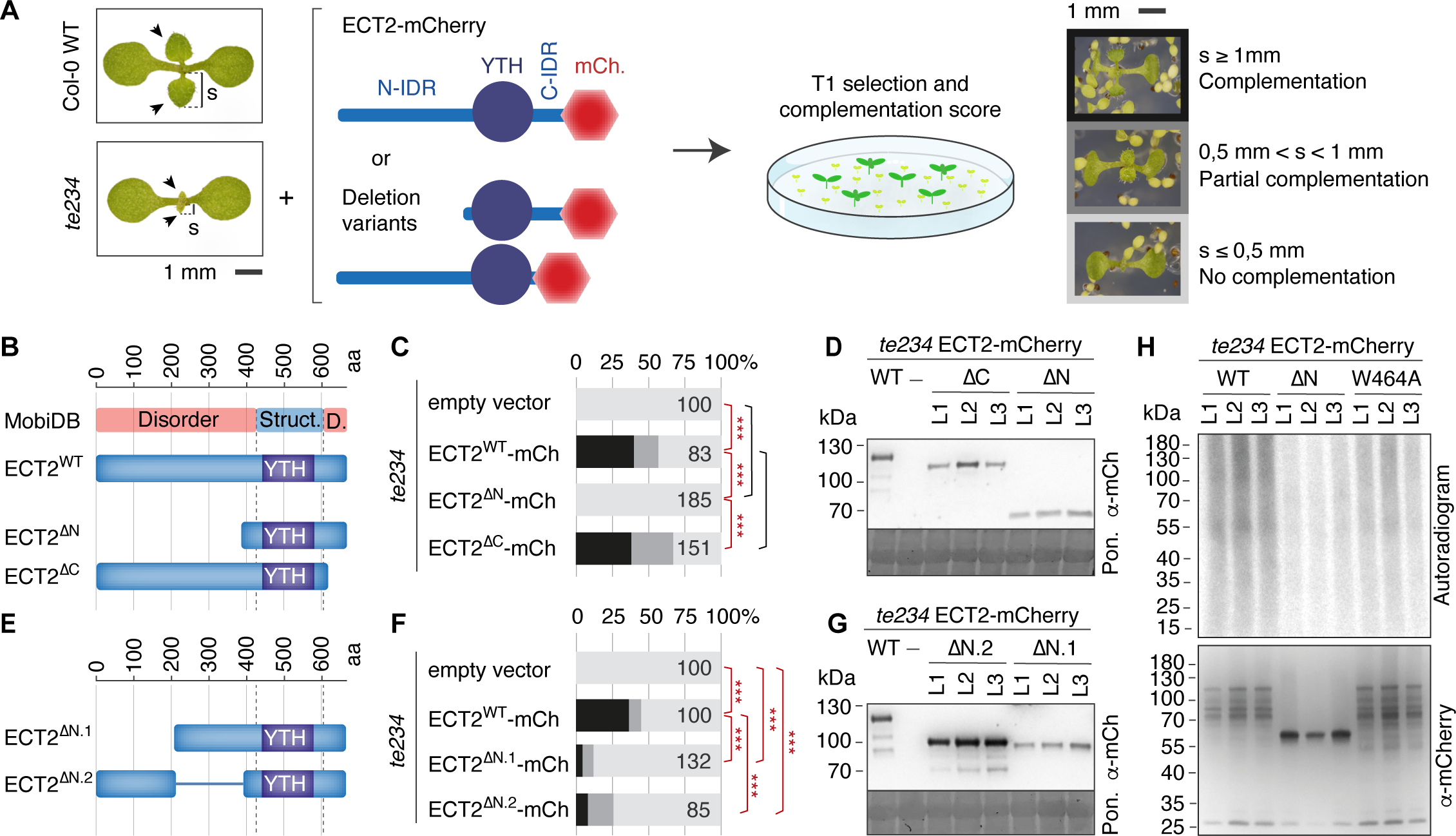
The long N-terminal IDR is indispensable for ECT2 function in plant development. A Graphical representation of the strategy followed for the assessment of the functionality of different regions of the ECT2 IDR. Left panel, seedlings of Col-0 WT or *te234* mutant showing the difference in the size (s) of the first true leaves (arrowheads) 9 days after germination. Next panels, the progeny of *te234* plants transformed with wild type or deletion variants of *ECT2p:ECT2-mCherry* are selected, and seedlings are classified into three categories according to s (right panel). B Schematic representation of wild type and mutant ECT2 proteins. The MobiDB (Di Domenico *et al*, 2012) track (top) shows the regions predicted to be structured (Struct.) or disordered (D.) along the protein length in amino acids (aa). Sequences of deletion mutants are listed in Data Set EV1. C Percentage of T1 seedlings falling into the three classes defined in A. The number of independent transformants analysed for each genotype (n) is indicated on the right side of the bars. Red lines with asterisks indicate significant differences according to pairwise Fisher exact test with Holm-adjusted *p*-values (* < 0.05, **< 0.01, *** < 0.001). Black lines indicate no significant difference. D Western blot showing the amount of ECT2-mCherry protein variants in T2 seedlings of three independent transgenic lines (L1-L3) per type. Ponceau (Pon) staining is used as loading control. E-G Same as B-D for the deletion mutants lacking halves of the N-terminal IDR. Corresponding protein sequences are listed in Data Set EV1. H Upper panel, autoradiogram showing PNK-labelled RNA co-purified with ECT2-mCherry variants after UV-crosslinking. Three independent lines for each ECT2-mCherry variant were analysed. Lower panel, western blot showing the amount of ECT2-mCherry variants from the immunopurifications above.

### The N-terminal, but not the C-terminal, IDR is required for function

We first designed and cloned mCherry-tagged *ECT2* transgenes devoid of either the first 410 amino acids (aa) in the large N-terminal IDR (*ECT2^ΔN^-mCherry*), or the last 56 aa in the shorter C-terminal IDR (*ECT2^ΔC^-mCherry*) (Fig 1B, Data Set EV1). After transformation of *te234* plants, functionality was measured as the complementation frequency in the first transgenic generation compared to the *ECT2^WT^-mCherry* construct for which about 60% complementation frequency is routinely obtained (Arribas-Hernández *et al*., 2018). T-DNA silencing is the main reason that complementation frequencies do not reach 100% with a wild type construct (Flores-Téllez *et al*., 2023). Expression of the wild type and deletion mutants in *te234* showed that ECT2^ΔC^ was fully functional while ECT2^ΔN^ lost function (Fig 1B-D). We further subdivided the 410-aa ΔN deletion into two ∼200-aa deletions, an N-terminal (ΔN1) and a C-terminal (ΔN2) half (Fig 1E, Data Set EV1). In both cases, the mutant proteins with only half IDR retained only residual activity (Fig 1F-G). Thus, the N-terminal IDR of ECT2, as well as its N-terminal and C-terminal halves individually, are essential for function *in vivo*.

### The N-terminal IDR is required for full RNA binding activity of ECT2 in vivo

We next used UV-crosslinking to assess whether the IDR is required for RNA association *in vivo.* We exposed entire seedlings to UV light, subjected ECT2^WT^-mCherry or ECT2^ΔN^-mCherry to immunoaffinity purification, and visualized the amount of crosslinked, immunoprecipitated (CLIPed) RNA by radiolabeling with polynucleotide kinase (PNK) after DNAse treatment (Arribas-Hernández *et al*., 2021a). We also included the ECT2^W464A^-mCherry mutant with a lesion in the m^6^A-binding aromatic cage as a control that exhibits strongly decreased, yet not abolished RNA association *in vivo* (Arribas-Hernández *et al*., 2021a). These experiments showed that RNA association was clearly reduced in ECT2^ΔN^ compared to wild type, to levels similar to those detected with the m^6^A-binding deficient mutant (Fig 1H). Thus, both IDR and YTH domains are necessary for full RNA-binding activity *in vivo*. This result has important implications for how specificity in YTHDF-target mRNA interactions may be achieved, and is consistent with our previous analysis that uncovered CLIP-seq tags attributable to both YTH-domain and IDR parts of ECT2 (Arribas-Hernández *et al*., 2021a). We note that the present result does not allow an assessment of whether the requirement of the IDR for RNA association relies on direct RNA contacts, interaction with other RNA binding proteins, or a combination of the two. We also note that the result does not exclude important functions of the N-terminal IDR other than those implicated in RNA association.

### Two distinct 30-40 aa regions in the N-terminal IDR, N3.2 and N8, are required for ECT2 function in vivo

Having established that the N-terminal IDR is required for ECT2 function, we next searched for smaller functional elements using a mutational approach. To design such mutants, we first analyzed sequence features of the N-terminal IDR with an eye towards the following two properties: (i) amino acid composition, as this gives clues to biophysical properties such as ability to engage in phase separation (Kato *et al*, 2012; Vernon *et al*, 2018), and (ii) conservation among ECT2 orthologues in dicotyledonous plants (Fig 2A). Based on these features, we designed six smaller deletions in the N-terminal IDR of ECT2 (ΔN3-ΔN8), each 40-70 aa in size (Fig 2A-B, Data Set EV1) and assessed their functionality as above. The results showed that despite protein accumulation similar to ECT2^WT^, ECT2^ΔN3^ and ECT2^ΔN8^ gave significantly lower complementation frequencies than wild type. In contrast, no significant reductions in complementation frequency were found for ECT2^ΔN4^-ECT2^ΔN7^ (Fig 2C-D), with ECT2^ΔN6^ even giving rise to slightly higher complementation frequencies than ECT2^WT^ (Fig 2D). We previously verified that ECT2^ΔN5^ fulfills developmental functions similar to wild type ECT2, but exhibits reduced antiviral activity (Martínez-Pérez *et al*., 2022), underscoring the fact that our functional complementation assay based on leaf emergence identifies elements required for developmental functions of ECT2, not necessarily all of its functions. We further subdivided the N3 element into two 40-aa segments (N3.1 and N3.2) (Fig 2A (bottom) and E, Data Set EV1) and found that ECT2^ΔN3.2^ was as defective as ECT2^ΔN3^, while ECT2^ΔN3.1^ retained wild type function (Fig 2C and 2F). We also combined ΔN3.2 and ΔN8 (Fig 2G) and observed a largely additive effect of the two deletions on complementation frequency (Fig 2H), suggesting that independent mechanisms of action underlie their requirement for function. We conclude that the deletion analysis identified two independently acting elements whose removal from the N-terminal IDR of ECT2 results in partial loss of function: Element N3.2 (G48-Y86) defined by the ΔN3.2 deletion, and element N8 (D355-K394) defined by the ΔN8 deletion. The remainder of this report focuses on understanding the properties of N3.2; similar work on N8 will be reported elsewhere.

**Figure 2.**
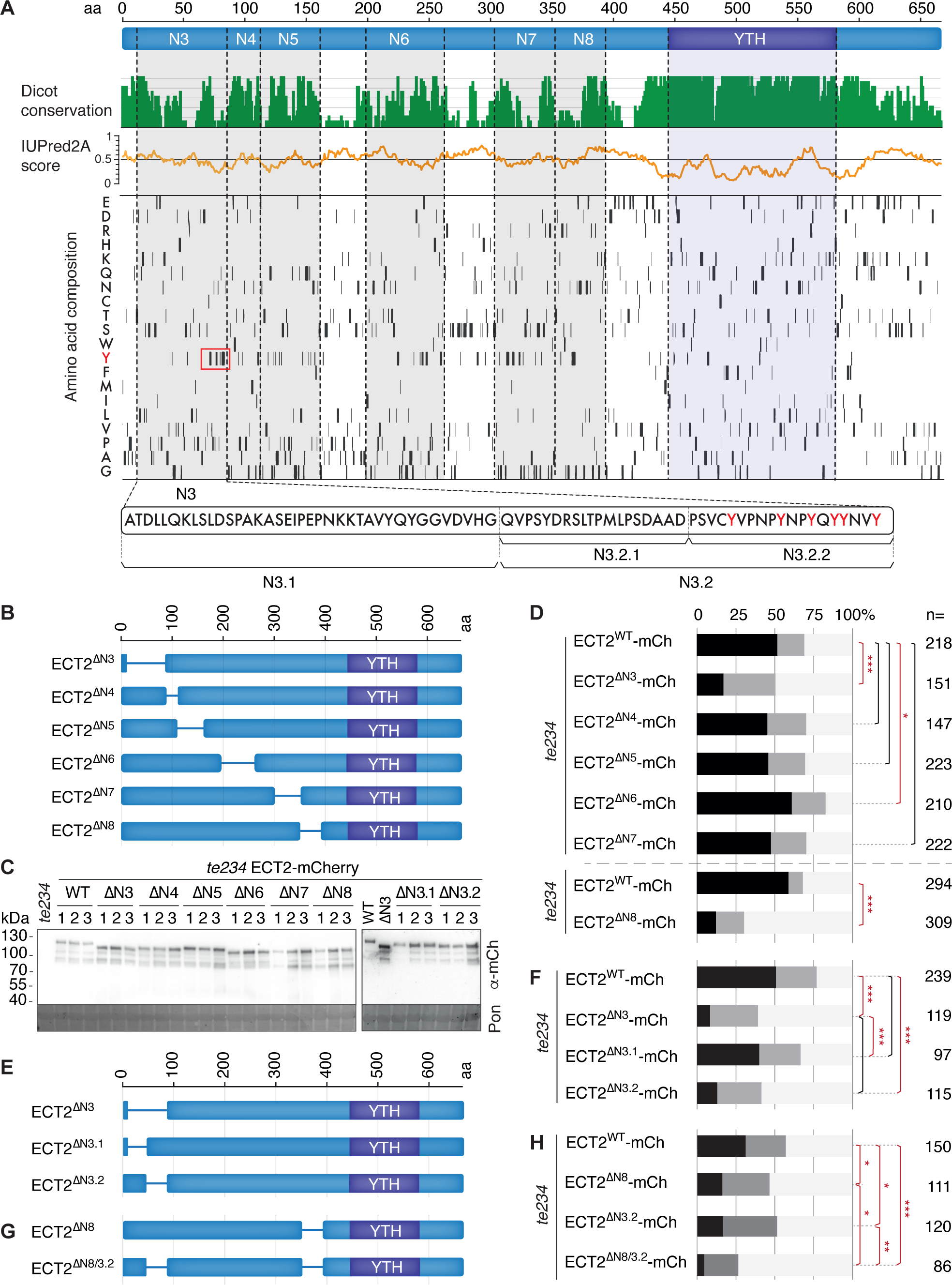
Two independently acting elements in the IDR are key for ECT2 function in plant development. A Analysis of ECT2 amino acid sequence. Top track: conservation between ECT2 orthologs in different dicotyledonous plant species. Second track: IUPred2A score [0-1] corresponding to the probability of the given residue being part of a disordered region (Mészáros *et al*, 2018). Bottom: amino acid composition. The N3-N8 elements in the N-terminal IDR selected for functional assays are indicated. Additional partitions of N3 are specified below the main panel, and the abundant tyrosine residues present in its C-terminal part (N3.2.2) are marked in red. B Graphical representation of the ΔN3-ΔN8 ECT2 deletion mutants. Corresponding protein sequences are listed in Data Set EV1. C Western blot showing the amount of ECT2-mCherry protein variants in T2 seedlings of three independent transgenic lines (1-3) per type. Ponceau (Pon) staining is used as loading control. D Complementation assay for ΔN3-ΔN8 ECT2 deletion mutants as in Fig 1C. Red lines with asterisks indicate significant differences according to pairwise Fisher exact test with Holm-adjusted *p*-values (* < 0.05, **< 0.01, *** < 0.001). Black lines indicate no significant difference. E Same as B for the two halves of the ΔN3 ECT2 deletion. Corresponding protein sequences are listed in Data Set EV1. F Same as in D for ECT2^ΔN3.1^ and ECT2^ΔN3.2^ compared to ECT2^ΔN3^ and wild type ECT2. G Same as B for the combined ΔN3.2 and ΔN3.8 ECT2 deletions. Corresponding protein sequences are listed in Data Set EV1. H Same as in D for ECT2^ΔN8/3.2^ compared to ECT2^ΔN3.2^, and ECT2^ΔN8^, and wild type ECT2.

### N3.2 is not required for RNA binding in vivo

We first tested whether RNA binding *in vivo* was compromised in the ECT2^ΔN3.2^ protein using PNK labeling of RNA co-immunoprecipitated with ECT2^ΔN3.2^-mCherry and ECT2^WT^-mCherry after UV-crosslinking *in vivo*. This assay did not show reduced PNK labelling of RNA co-purified with ECT2^ΔN3.2^ compared to ECT2 wild type. Rather, labelling of ECT2^ΔN3.2^-RNA complexes was more intense than of ECT2^WT^-RNA complexes (Figs 3A and EV1A). Because ECT2-RNA complexes with or without the N-terminal IDR generated by proteolysis during immuno-purification are labeled with different efficiency, presumably due to different accessibility of 5’-ends as a consequence of IDR-RNA contacts (Arribas-Hernández *et al*., 2021a), we think it unwise to interpret this result more deeply than to conclude that ECT2^ΔN3.2^ retains RNA-binding capacity. We also note that this experiment does not provide insight into the identities of the co-purified mRNAs, and it remains possible that N3.2 is implicated in mRNA target selection. Nonetheless, taken together with the fact that ECT2/3/4 are major mediators of developmental m^6^A functions (Arribas-Hernandez & Brodersen, 2020), the indication of intact RNA-binding activity of ECT2^ΔN3.2^ suggests that m^6^A is unlikely to act only via protection against local endonucleolysis, as proposed previously (Anderson *et al*., 2018). This indication also led us to focus our attention on properties other than RNA binding to understand the function of N3.2.

**Figure 3.**
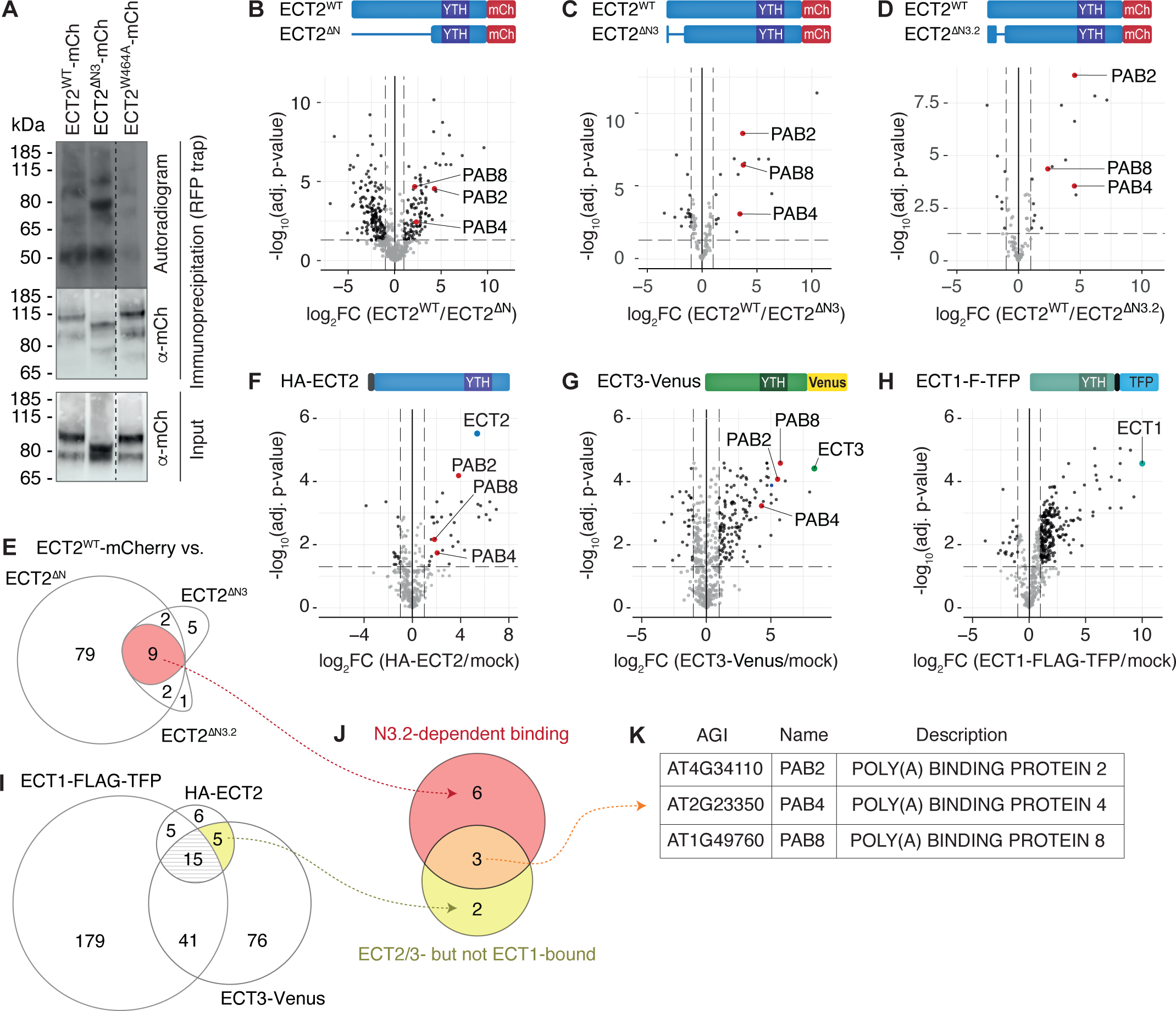
The N3 element is necessary for interaction between ECT2 and PAB2/4/8. A CLIP analysis of the amount of RNA immunopurified with ECT2-mCherry wild type and mutant variants (Δ3 and the aromatic cage mutant W464A) as in Fig 1H. Tissue from three independent lines per genotype were pooled for the analysis in this case. B-D Volcano plots showing the differential abundance of proteins co-purified with variants of ECT2-mCherry (RFP-trap) as detected by mass spectrometry of immunopurified fractions (IP-MS). Diagrams with the proteins compared in each case are shown above. The poly(A)-binding proteins PAB2, 4 and 8 are specifically lost by deletions of the N3 element of the N-terminal IDR. E Overlap between significantly differentially abundant proteins (FC > 2, adjusted p-value < 0.05) found in the IP-MS experiments in B-D. F-H Same as in B-D for the proteins more abundant in HA-ECT2 (HA beads, F), ECT3-Venus (GFP-trap, G) and ECT1-FLAG-TFP (FLAG beads, H), purifications compared to a mock IP with non-transgenic plants. I Same as in E for the IP-MS experiments shown in F-H. J Overlap showing N3-dependent ECT2 interactors also found in HA-ECT2 and ECT3-GFP purifications, but not in the ECT1-FLAG-TFP fractions. K List of the three only genes found in the overlap in J.

### The N-terminal IDR element N3.2 of ECT2 is required for interaction with the cytoplasmic PABPs, PAB2/4/8

We next tested the possible importance of N3.2 in protein-protein interaction. We conducted immunoprecipitation-mass spectrometry (IP-MS) analyses of RNAse-treated lysates prepared from plants expressing mCherry fusions of either ECT2^WT^, ECT2^ΔN^, ECT2^ΔN3^, or ECT2^ΔN3.2^ (Fig EV1B-C). These analyses showed that while the full N-terminal IDR was required for association with almost a hundred proteins *in vivo* (Fig 3B and Data Set EV2), only 16 and 12 proteins showed significant depletion in ECT2^ΔN3^ and ECT2^ΔN3.2^ purifications, respectively, compared to ECT2^WT^, and 9 of them were common between the three datasets (Fig 3C-E and Data Set EV3). To further validate and narrow the number of interactors of possible functional significance, we completed four additional IP-MS analyses using hemagglutinin (HA)-tagged ECT2, Venus-tagged ECT3, and FLAG-TFP tagged ECT1 (Arribas-Hernández *et al*., 2018; Flores-Téllez *et al*., 2023) (Figs 3F-H and EV1D-F, and Data Set EV2). Because ECT2 and ECT3 exhibit genetic redundancy in plant development (Arribas-Hernández *et al*., 2018; Arribas-Hernandez *et al*., 2020), we reasoned that interactors necessary for ECT2 activity must co-purify with both proteins. On the other hand, ECT1 does not have the developmental function of ECT2/3 despite its strong similarity to ECT3 (55% amino acid identity), and therefore such interactors may be absent from ECT1 purifications. The comparison between the proteins identified in each purification compared to mock IPs from non-transgenic plants showed that 20 candidates had a significant enrichment in HA-ECT2^WT^ and ECT3-Venus fractions, and that 5 of them were not found in ECT1 pulldowns (Fig 3I and Data Set EV3). Remarkably, 3 of these proteins, the poly(A) binding proteins PAB2, PAB4 and PAB8, were also among the 9 N3.2-dependent interactors (Fig 3J-K), and are, therefore, outstanding candidates for functionally important interactors. Poly(A) binding proteins associate with the poly(A) tail of mRNA to effect a multitude of functions related to mRNA translation, and PAB2, PAB4 and PAB8 are the three most highly and broadly expressed paralogs among the eight cytoplasmic poly(A)-binding proteins (PABPs) in arabidopsis (Belostotsky, 2003; Goss & Kleiman, 2013). Thus, we proceeded to characterize ECT2-PABP interactions in more detail.

### ECT2 function and PABP interaction relies on a small Tyr-rich region

To reinforce the link between ECT2 function and PABP interaction, we further subdivided ΔN3.2 into two halves (Figs 2A (bottom) and 4A, Data Set EV1) and used these smaller deletion mutants in the *te234* complementation assay (Fig 1A). This effort narrowed the functionally important part to the region defined by ΔN3.2.2, a 19-amino acid region containing six tyrosine residues (Figs 2A and 4A-B). These six tyrosine residues are required for function, because their mutation into alanine (ECT2^6xY➔A^) resulted in loss of ECT2 function comparable to that caused by the ECT2^ΔN3^ deletion mutant, despite protein accumulation comparable to ECT^WT^ (Figs 4C-D and EV2, Data Set EV1). We finally used the refined ECT2^ΔN3.2.2^ mutant for IP-MS analysis that also showed loss of PAB2, PAB4 and PAB8 interaction *in vivo* (Fig 4E). Thus, loss of ECT2 function in mutants in the N3 region correlates closely with the loss of association with PAB2, PAB4 and PAB8, suggesting that the physical association between ECT2 and cytoplasmic PABPs is functionally important. ECTs and PAB2/4/8 have previously been found to co-purify with stress granule markers even in the absence of stress (Kosmacz *et al*, 2019), and a recent report also found an association between ECT2 and PAB2 to be mediated by a 400-aa N-terminal IDR fragment of ECT2 (Song *et al*, 2023). Although two 200-aa fragments corresponding to N- and C-terminal halves of the IDR were sufficient for PAB2 interaction in qualitative *in vitro* binding assays, the region necessary for interaction was not more accurately defined (Song *et al*., 2023).

**Figure 4.**
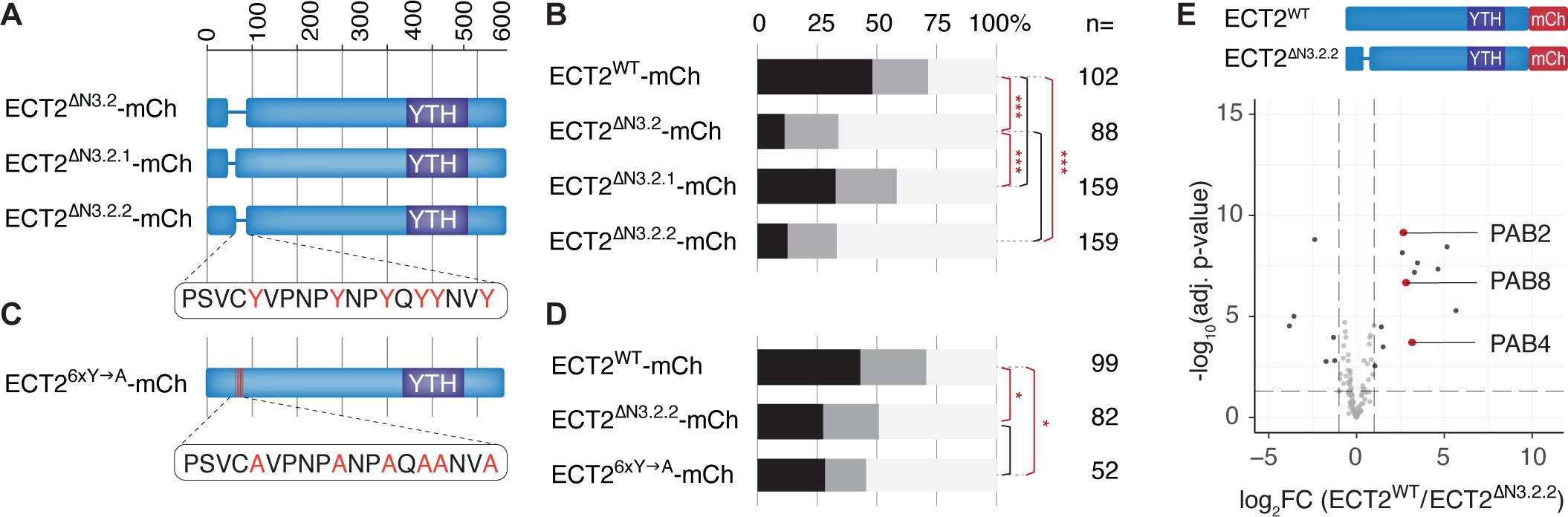
The tyrosine-rich region in N3 is essential for ECT2 function. A Graphical representation of the partition of ΔN3.2 into ΔN3.2.1 and ΔN3.2.2. Corresponding protein sequences are listed in Data Set EV1. B Complementation assay of ΔN3.2.1 and ΔN3.2.2 ECT2 deletion mutants as in Fig 1C. Red lines with asterisks indicate significant differences according to pairwise Fisher exact test with Holm-adjusted *p*-values (* < 0.05, **< 0.01, *** < 0.001). Black lines indicate no significant difference. C Graphical representation of the mutations in ECT2^6xY➔A^. The exact corresponding protein sequence is given in Data Set EV1. D Complementation assay ECT2^6xY➔A^ as in B. E IP-MS analysis of the ECT2-interacting proteins lost upon deletion of N3.2.2 compared to ECT2^WT^ as in Fig 3B-D.

### Increased PAB4 dosage is sufficient to suppress leaf formation defects in te234 mutants

We next sought genetic evidence to corroborate the functional importance of the ECT-PABP interaction. Since PAB2, PAB4 and PAB8 are largely functionally redundant in strict genetic terms during vegetative growth, and *pab2 pab4 pab8* triple mutants are embryonically lethal (Dufresne *et al*, 2008; Gallie, 2017; Zhao *et al*, 2019), we could not employ standard knockout analyses. Nonetheless, physically interacting components of biological systems often show dosage dependence, such that the importance of their interaction can be revealed genetically through either non-allelic non-complementation (Kidd *et al*, 1999) or overexpression suppression (van Leeuwen *et al*, 2017). Given the number of loci involved (*ECT2/ECT3/ECT4/PAB2/PAB4/PAB8*) and the clear phenotype of *te234* mutants, the most straightforward genetic test is the effect of PABP overexpression in organ primordia of *te234* mutants. Thus, we expressed C-terminally Venus-tagged PAB4 in *te234* mutants under the control of the strong *US7Y/RPS5A* promoter specific for rapidly dividing cells (Weijers *et al*., 2001) (Fig 5A). After 9 days of growth, a clear rescue of delayed leaf emergence was visible in *PAB4-Venus*-expressing *te234* mutants compared to *te234* seedlings without the *PAB4* transgene (Fig 5B-D), and partial suppression of the defects in the size and shape of the leaves could be appreciated at later stages (Fig 5B). Thus, increased PAB4 dosage is sufficient to alleviate the delayed leaf formation resulting from the simultaneous knockout of *ECT2*, *ECT3* and *ECT4*, supporting the functional relevance of the interaction between these essential families of RNA-binding proteins.

**Figure 5.**
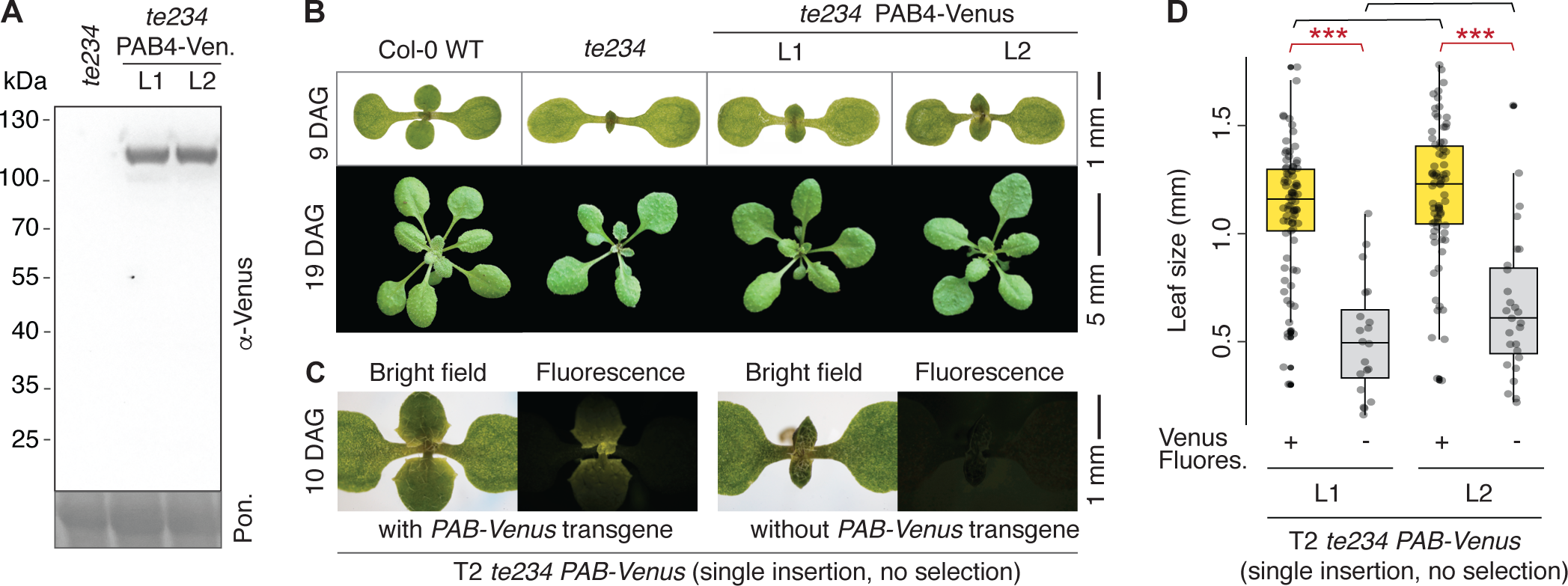
Overexpression of PAB4 partially suppresses defective leaf formation caused by loss of ECT2/3/4 function. A Western blot analysis showing expression of the *US7Yp:PAB4-Venus-OCSt* (*PAB4-Venus*) transgene in *te234* plants. Notice the integrity of the fusion protein. Ponceau (Pon.) staining is used as loading control. B Partial suppression of the delay in leaf formation (top) and defective leaf morphology (bottom) of *te234* plants in two independent transgenic lines (L1, L2) overexpressing PAB-Venus in actively dividing cells. DAG, days after germination. C Morphological appearance and Venus fluorescence in the first true leaves of a 10-day-old T2 seedling expressing PAB4-Venus in the *te234* background (left) compared to a sibling without PAB4-Venus expression (right) due to segregation of the single-insertion transgene in T2. D Box plot showing the quantification of the size of leaves of 10-day-old T2 seedlings of two independent *te234 PAB4-Venus* lines. Because they have single T-DNA insertions, the transgene segregates in the T2 population and the leaf size of sibling plants with or without the transgene (Venus + or – respectively) growing side by side can be measured. Significance was assessed using Student’s *t*-test (\*\*\**p* < 0.0001).

We note that PAB2, PAB4 and PAB8 mRNAs are all high-confidence m^6^A-ECT2/3 targets (Arribas-Hernández *et al*., 2021b; Parker *et al*., 2020; Shen *et al*., 2016) (Fig EV3A). It is, therefore, possible that their expression may be lower in *ect2 ect3 ect4* mutants than in wild type, and that such a difference in expression might contribute to the slow leaf formation and explain the partial phenotypic rescue by PAB4 overexpression. We consulted our previously published transcriptome analyses to address this possibility (Arribas-Hernández *et al*., 2021b). In entire root tips, no significant differences in PAB2/4/8 mRNA expression were observed between *ect2 ect3 ect4* mutants and wild type (Fig EV3B). However, in *ECT2*-expressing root protoplasts selected by fluorescence-assisted cell sorting, a ∼3-fold downregulation of *PAB4* and slightly less than 2-fold downregulation of *PAB8* mRNAs were in fact observed in cells devoid of ECT2/3/4 function compared to *ECT2*-expressing controls (Fig EV3B). Thus, we cannot entirely exclude the possibility that simple downregulation of *PAB4* mRNA in *te234* mutants underlies the suppression of slow leaf formation by *PAB4* overexpression. We disfavor this possibility, because all single homozygous knockout mutants in *PAB2/4/8* genes have normal developmental phenotypes (Gallie, 2017; Zhao *et al*., 2019), suggesting that PABP function *in vivo* is not excessively sensitive to dosage. In addition, the argument in favor of functional importance of the physical association of ECT2 and PAB2/4/8 relies not only on the suppression of *te234* mutant phenotypes by PAB4 overexpression in organ primordia, but also on the tight correlation between loss of ECT2 function and loss of PAB2/4/8 association.

Based on the ECT2-PAB2/4 interaction, and the observation that some ECT2 mRNA targets have shorter half-lives in the absence of ECT2/3/4, it was recently proposed that m^6^A-ECT2/3/4 generally acts to stabilize mRNA targets via PAB2/4 interaction (Song *et al*., 2023). This proposition is not supported by evidence, however, because no efforts were made to specifically disable the ECT2-PAB2/4 interaction, or to restore PAB2/4 binding to ECT2/3/4 mRNA targets in the absence of m^6^A or ECT2/3/4. Even with the clear correlation between partial loss of ECT2 function and PAB2/4/8 interaction in ECT2^ΔN3^-mutants, and the suppression of the *ect2 ect3 ect4* phenotype by PAB4 overexpression shown here, we propose a rather less bold interpretation of the evidence at hand. We suggest that the concentration of PAB2/PAB4/PAB8 in rapidly dividing primordial cells is limiting, and that at least a subset of ECT2/3/4 target mRNAs requires the ECT-PABP interaction to compete efficiently for PABP binding with other mRNAs. Increasing the cytoplasmic PABP concentration by PAB4 overexpression would, therefore, alleviate the requirement for m^6^A-ECT2/3/4 to ensure sufficient PABP binding to this subset of target mRNAs. Because in early vertebrate embryogenesis, mRNAs with short poly(A) tails were recently shown to compete less efficiently for limiting amounts of PABP than those with longer poly(A) tails (Xiang & Bartel, 2021), we also suggest that the ECT2/3/4 targets relevant for organogenesis are likely to be found among mRNAs with short poly(A) tails. We also note that a requirement for cytoplasmic PABP for viral translation and infectivity, as demonstrated in the case of Turnip Mosaic Virus (Dufresne *et al*., 2008), may be part of the explanation why metabolically active, primordial cells would evolve to function with limiting PABP concentrations and a requirement for some mRNAs to use the m^6^A-ECT axis to compete efficiently for PABP binding. Nonetheless, other models to explain the importance of ECT2-PAB interaction cannot be excluded at this point. For example, rather than stabilizing PAB binding to m^6^A-ECT2/3 target mRNAs, ECT-PAB interaction might increase the rate of exchange of nuclear and cytoplasmic PABP (Mangus *et al*, 2003) to accelerate formation of translation-competent mRNPs upon import into the cytoplasm.

## MATERIALS AND METHODS

### Plant material and growth conditions

All the plant lines used in this study are in the *A. thaliana* Columbia-0 (Col-0) ecotype. The following mutant and transgenic lines have been previously described: *ect2-1 ect3-1 ect4-2* (*te234*), *ect3-1 ect4-2* (*de34*)*, ect2-1 ECT2-mCherry*, *ect2-1 ECT2^W464A^-mCherry, ect2-1 HA-ECT2*, *ect3-2 ECT3-Venus*, *ect4-2 ECT4-Venus* (Arribas-Hernández *et al*., 2018), and *ect1-2 ECT1-FLAG-TFP* (Flores-Téllez *et al*., 2023). Seeds were sterilized by immersion in 70% EtOH for 2 min followed by incubation in 1.5% NaOCl, 0.05% Tween-20 for 10 min, and immediately washed twice H_2_O. The seeds were spread on plates containing Murashige & Skoog (MS) medium (4.1 g/L MS salt, 10 g/L sucrose, 8 g/L Bacto agar). Plates were stratified in darkness at 4°C for 2-5 days before transfer to Aralab incubators at 21°C, with a light intensity of 120 µmol/m^2^ and 16 h light / 8 h dark photoperiod. Seedlings for UV-crosslinking and IP were grown on vertical plates. When necessary, seedlings were transferred to soil after 10 days of *in vitro* growth and maintained in Percival incubators with identical settings.

### Construction of ECT2 deletion mutants and PAB4 overexpressors

To introduce deletions or point mutations in *ECT2p:ECT2-mCherry-ECT2t,* and to generate *US7Yp:PAB4-Venus-OCSt* transgenes, we employed the scar-free USER cloning method (Bitinaite and Nichols, 2009) to piece together PCR-amplified DNA fragments in all cases except for the 6xTyr-to-Ala mutation, for which appropriate dsDNA (Data Set EV4) for USER cloning was directly synthesized (Integrated DNA Technologies, gBlocks). As template for PCR, we used plasmids containing wild type *ECT2p:ECT2-mCherry* (genomic DNA) (Arribas-Hernández *et al*., 2018) for *ECT2-mCherry* constructs, GreenGate pGGA012 for the *US7Y/RPS5A* promoter, GreenGate pGGF005 for the *OCS* terminator (Lampropoulos *et al*, 2013), and *ECT3p:ECT3-Venus* (Arribas-Hernández *et al*., 2018) for *Venus*. The genomic sequence of PAB4 (At2g23350) was amplified from Col-0 WT genomic DNA, prepared as previously described (Arribas-Hernandez *et al*, 2016). The fragments were amplified with dU-substituted primers and KAPA Hifi Hotstart Uracil+ ReadyMix (Bitinaite & Nichols, 2009) and inserted in the pCAMBIA3300-U vector (pCAMBIA3300 with a double PacI USER cassette inserted between the PstI-XmaI sites at the multiple cloning site) (Nour-Eldin *et al*, 2006). The resulting plasmids were checked by restriction digestion analysis and sequencing before their transformation into plants via *Agrobacterium-*mediated floral dip (Clough & Bent, 1998) using the GV3101 strain (Koncz & Schell, 1986). The sequences of primers used for every fragment of each construct are detailed in Data Set EV4.

### Screening for te234 complementation

Primary transformants (T1s) of *te234* carrying wild type, deletion or point mutation variants of *ECT2-mCherry* were selected on 15-cm diameter MS-agar plates supplemented with glufosinate ammonium 7.5 mg/L (Sigma) to select plants with the transgene, and ampicillin 10 mg/L to restrain the growth of agrobacteria. Nine days after germination, each primary transformant was binned into one of three categories according to the size (s) of the first true leaves: full complementation (s IZ 1 mm), partial complementation (0.5 mm < s < 1 mm), or no complementation (s IZ 0.5 mm). The complementation percentages were then calculated as the number of seedlings in each complementation category relative to the total number of transformants.

### Statistical analyses

Fisher’s exact test was employed to determine the statistical significance of the observed differences among the T1 complementation categories and the Holm-Bonferroni method was used to account for multiple testing. Student’s *t*-test was used to assess the significance of difference in leaf size between *te234* mutants carrying or not the *US7Yp:PAB4-Venus-OCSt* transgene.

### ECT2-IDR analyses

To assess conservation among dicot ECT2 homologs we performed a protein alignment using CLUSTAW and transformed it into a conservation score (Livingstone and Barton, 1993). The geneIDs used for the alignment are the following; *Helianthus annuus* (A0A251VJQ5), *Lupinus angustifolius* (LOC109348290), *Vitis vinifera* (A5C6V5), *Solanum tuberosum* (M1C7V3), *Theobroma cacao* (A0A061GSW6), *Beta vulgaris subsp*. *vulgaris* (LOC104888716), *Corchorus capsularis* (A0A1R3IEW9), *Brassica rapa* (LOC103859518), *Phaseolus angularis* (A0A0L9TJY9), *Medicago truncatula* (G7L842), *Daucus carota subsp. sativus* (A0A164SV78), *Manihot esculenta* (A0A2C9VJN4), *Phaseolus vulgaris* (V7CQK6), *Glycine max* (I1KSN3), *Populus trichocarpa* (B9GXX7), *Gossypium raimondii* (A0A0D2P5E4), *Trifolium pratense* (A0A2K3P0Z0). The prediction of protein disorder was performed utilizing the publicly available webserver IUPred2A in default mode (Mészáros *et al*, 2018).

### Western blotting

100 – 300 mg of powdered tissue was mixed with 5x (v/w) IP buffer IP buffer (50 mM Tris-HCl ph 7.5, 150 mM NaCl, 10% glycerol, 5 mM MgCl2, and 0.1% Nonidet P40) freshly supplemented with protease inhibitor (Roche Complete tablets) and 1 mM DTT (reducing agent). The lysate was mixed immediately by vigorous shaking and spun at 13.000 g for 10 minutes. 4x LDS sample buffer (277.8 mM Tris-HCl pH 6.8, 44.4% (v/v) glycerol, 4.4% LDS, and 0.02% bromophenol blue) was mixed with the protein samples to a concentration of 1x LDS. The samples were denatured at 75° C for 10 minutes before running on a 4-20% Criterion^TM^ TGX^TM^ Precast gel and run in 1x Tris Glycine, 0.1% SDS at 90-120 V for ∼1 hour on ice. Using wet transfer, proteins were immobilized by blotting them onto an Amersham Protran Premium nitrocellulose membrane (GE Healthcare Life Sciences) in cold 1 x Tris-Glycine, 20% EtOH transfer buffer at 80 V for 1 hr on ice. Following transfer, the membrane was blocked in 5 % skimmed milk, 0.05% Tween-20, PBS buffer (137 mM NaCl, 2.7 mM KCI, 10 mM Na_2_HPO_4_, 1.8 mM KH_2_PO_4_, pH 7.4) for 30 min. After blocking, membranes were incubated with commercially available antibodies against mCherry (Abcam ab183628,1:2000 dilution) or Venus (Sicgen AB2166-100; 1:1000 dilution). The following day, membranes were washed in PBS-T (PBS buffer with Tween-20 (Bio-Rad) to a total concentration of 0.05%).

### Immunoprecipitation for mass spectrometry

To immunoprecipitate ECT proteins, ∼300 mg of liquid nitrogen-ground tissue powder from 9-day old seedlings were added to 1 mL of IP buffer (50 mM Tris-HCl ph 7.5, 150 mM NaCl, 10% glycerol, 5 mM MgCl2, and 0.1% Nonidet P40) with the following additives: 5 mM DTT, 2 mM AEBSF (Sigma), 40 µg/mL RNase A, and 1.5 tablet/10 mL of cOmplete protease inhibitor. Sigma plant protease inhibitor (Sigma P9599) was also added to IPs (200 µL/10 mL of IP buffer) of *ect2-1* HA-ECT2 and *ect2-1* ECT2-mCherry. Immediately after vigorous vortexing, the lysates were centrifuged at 4°C for 10 minutes (∼13.000 *g*), and the supernatants were passed through hydrophilic 0.45 µm filters for incubation with 12.5 µL of washed beads (RFP-TRAP®_A (Chromotek) for all ECT2-mCherry variants, GFP-TRAP®_A (Chromotek) for ECT3-Venus, Anti-HA affinity Matrix (Roche) for HA-ECT2 and ANTI-FLAG M2 affinity resin (Sigma) for ECT1-FLAG TFP). After 1 hour of slow rotation at 4°C, the beads were washed three times (5 min each) with ice-cold IP-buffer supplemented only with 4 mM DTT. To remove detergent and excess salt, we performed two additional washes with ice-cold 1xPBS. After the final wash, the purifications were either eluted using 100 ug/mL FLAG peptide (ECT1-FLAG-TFP) or the supernatant was removed from the beads (all ECT2-mCherry variants, ECT3-Venus, HA-ECT2). In any case, samples were kept at −80C until tryptic digestion for mass spectrometry. The purifications were done in three replicates for both the transgenic lines expressing tagged ECTs, or non-transgenic plants for mock IPs.

### Mass spectrometry

Purifications were prepared for mass spectrometry by trypsin/LysC digestion and C18 stage tipping. Beads containing purified proteins were resuspended in 20 µL lysis buffer (6M guanidium chloride, 10 mM Tris(2-carboxyethyl)phosphine, 40 mM 2-chloroacetamide (CAA), 50 mM HEPES pH 8.5). Samples were sonicated with three 30’’ on – 30’’ off cycles using BioRupter Pico and diluted 1:3 by adding 40 µL of digestion buffer (10 % acetonitrile, 50 mM HEPES pH 8.5). 2 µL of LysC (50 ng/ µL) were then added to the samples and incubated at 37° C for 3-4 hours, while shaking at ∼1500 rpm. Subsequently, samples were diluted 1:10 by adding 140 µL more of digestion buffer and 1 µL of trypsin (50 ng/ µL) was added, then briefly vortexed, and incubated at 37°C over night while shaking at ∼1500 rpm. The reaction was stopped by adding 2% trifluoro acetic acid (1:1, 1% final concentration), ie. 200 µL after which samples were vortexed and spun down for 1’ at 4000x rpm. 200 µL pipette tips were packed with two C18 filters (Empore 2215. 66883-U). The C18 filters were activated by adding; 30 µL MeOH, 30 µL Buffer B (80% acetonitrile, 0.1% Formic Acid), 2x 30 µL Buffer A’ (3% acetonitrile, 1% trifluoracetic acid). At this point, samples were loaded onto the StageTip with 50 µL aliquots at a time and spun through at ∼3,000x rpm. The tips were then washed twice with 100 µL of Buffer A (0.1 % FA) and eluted with 2x 30 µL Buffer B’ (40% I, 0.1 % FA) into a clean 500 µL Protein LoBind E. tube. The eluted samples were speedvaced for ∼60 mins at 60° C and resuspended in 12 µL Buffer A* (2 % I 1%TFA) containing iRT peptides. Finally, 1.5 µL were used for concentration measurements by Nanodrop. Non-targeted mass spectrometry analysis was performed on a quadrupole Orbitrap benchtop mass spectrometer, Qexactive (Thermo Scientific), equipped with an Easy nano-LC 1000 system (ThermoFisher Scientific). 500ng of peptide mixture from each sample was analyzed by online nano-scale liquid chromatography tandem mass spectrometry (LC-MS/MS) in turn. Peptides were separated on a 50 cm C18-column (Thermo EasySpray ES904 / Thermo EasySpray ES903) using an EASY-nLC 1200 system (Thermo Scientific). The column temperature was maintained at 45 °C. Buffer A consisted of 0.1% Formic acid in water, and buffer B of 80% CAN, 0.1% Formic acid. The flow rate of the gradient was kept at 250 nl/min, and started at 10% Buffer B, going to 23% buffer B in 85 minutes. This was followed by a 30’ step going to 38% buffer B, increasing to 60% Buffer B in 10 minutes, and finally ramping up to 95% buffer B in 5 minutes, holding it for 10 min. to wash the column. The Q Exactive Classic instrument (Thermo Scientific, Bremen, Germany) was run in data dependent acquisition mode using a top 10 Higher-energy Collisional Dissociation (HCD)-MS/MS method with the following settings. Scan range was limited to 350-1750 m/z. Full scan resolution was set to 70,000 m/z, with an AGC target of 3e6 and a maximum injection time (IT) value of 20ms. Peptides were fragmented with a normalized collision energy of 25, having a dynamic exclusion of 60s, excluding unassigned ions and those with a charge state of 1. MS/MS resolution was set at 17,500 m/z, with an AGC target of 1e6 and a maximum IT of 60ms.

### Analysis of IP-MS data

All raw LC-MS/MS data files were processed together using Proteome Discoverer version 2.4 (Thermo) with the use of Label-free quantitation (LFQ) in both the processing and consensus steps. In the processing step, Oxidation (M), and protein N-termini acetylation and met-loss were set as dynamic modifications, with cysteine carbamidomethyl set as static modification. All results were filtered with percolator using a 1% false discovery rate (FDR), and Minora Feature Detector was used for quantitation. SequestHT was used as database, matching spectra against the TAIR database. Two different methods of normalization were employed to infer the differentially enriched proteins between samples. When comparing ECT2-mCherry mutants to full length ECT2-mCherry, we made use of total sum normalization in addition to variance stabilizing normalization (VSN) (Huber *et al*, 2002). However, when analysing the differential abundance of proteins co-purified with ECTs compared to non-transgenic plants (mock IP), we decided to do only VSN normalization to prevent inflation of the background in the non-transgenic mock samples that contain very little protein overall. The VSN normalization method, implemented using the ‘normalizeVSN’ function from the limma package, was employed to transform the abundance values and stabilize the variance across the samples. To identify differentially abundant proteins between samples we made use of the R package limma (Smyth, 2004). First, a linear model was constructed using a design matrix and a contrast matrix to specify the contrasts between samples. Fold-changes and *p*-values were then calculated based on the linear model. We utilized the fitted linear model as an input to treat and topTreat which performed moderated t-statistics and log-odds estimation using Empirical Bayes statistics. *p*-values were adjusted for multiple testing using the Benjamini-Hochberg adjustment method (Benjamini & Hochberg, 1995).

### Silver staining

10-20% of the samples containing immunoprecipitated ECTs for mass-spectrometry were subjected to SDS-PAGE in 4-20% Criterion^TM^ TGX^TM^ Precast Midi Protein gels (Bio-Rad) for visual assessment. After electrophoresis at 90V, the gels were fixed with a solution of 30% EtOH,10% AcOH for 20 minutes and subsequently rinsed in a stepwise manner: 10 min in 20% EtOH, 10 min in 10% EtOH, and 2x 20 min in H_2_O. The gels were then submerged in sensitizing solution (0.02 M Na_2_S_2_O_3_) for 1 minute, rinsed two times with water, and incubated in 0.1% AgNO_3_ for 1 hour. After two additional washes with water, the gels were developed with freshly made developing solution (0,04% formalin, 2% Na_2_CO_3_) until the amount of signal was adequate, at which point the development was quenched with 2% AcOH.

### Analysis of RNA association in vivo by crosslinking-immunoprecipitation-PNK labeling

Crosslink, immunoprecipitation and PNK-labelling of ECT2-mCherry mutants were performed as previously described (Arribas-Hernández *et al*., 2021a). Briefly, 12-day-old seedlings were irradiated with 2000 mJ/cm^2^ of 254 nm-UV light on ice. The tissue was then ground in liquid nitrogen, and lysates containing ECT2-mCherry RNPs were prepared by resuspending 1 g of ground tissue in 1.5 mL of iCLIP buffer (0.25 % sodium deoxycholate, 0.25 Igepal, 50 mM Tris-HCL pH 7.5, 150 mM NaCl, 1% SDS, 5 mM DTT, 4 mM MgCl_2_) supplemented with protease inhibitors (Roche cOmplete 1 tablet/10 mL, 4 mM PMSF and 1/30 (v/v) Sigma Plant Protease Inhibitor). After homogenization, the lysates were cleared by centrifugation for 10 min at max. speed, passed through a 0.45 µm filter and incubated with 20 µL RFP-TRAP®_A (Chromotek) beads for 1 hour at 4°C under slow rotation. After removal of the supernatant, the beads were washed four times with iCLIP wash buffer (0.5 % sodium deoxycholate, 2M urea, 0.5% Igepal, 1% SDS, 50 mM Tris-HCL pH 7.5, 500 mM NaCl, 2 mM DTT, 4 mM MgCl_2_) and two times with PNK wash buffer (20 mM Tris-HCL, 10 mM MgCl_2_, 0.2% Tween-20). Beads were then resuspended in 100 µl PNK wash buffer and subjected to Dnase and Rnase digestion by addition of 2 µL Turbo Dnase (Thermo Fisher) and 5 µL of Rnase I (Ambion) (1:5000 dilution) for 10 min at 37°C and 1100 rpm. Beads were then washed once with PNK wash buffer, twice with high-salt buffer (50 mM Tris-HCl pH 7.4, 1 M NaCl, 1 mM EDTA, 1 % Igepal, 0.1 % SDS, 0.5 % sodium deoxycholate) and twice again with PNK wash buffer. The RNA crosslinked to ECT2 on the beads was radioactively labelled at the 5′ end by PNK-mediated phosphorylation using γ-^32^P-ATP (20’ at 37°C). After three washes in PNK wash buffer, beads were resuspended in 2x LDS buffer (Thermo Fisher), incubated at 95°C for 10’ and the denatured RNP complexes were then subjected to SDS-PAGE, blotted onto a nitrocellulose membrane (Protran BA-85) and detected by autoradiography.

### Fluorescence microscopy

Whole seedings and leaves were imaged employing a Leica MZ16 F stereomicroscope equipped with a Sony a6000 camera.

## Supporting information

Data Set EV1

Data Set EV2

Data Set EV3

## ACKNOWLEDGEMENTS

We thank Lena Bjørn Johansson, Daniel Tobias Kyndesen Lahti, Ida Thorøe Michler and Magnus von Holstein-Rathlou for technical assistance. Theo Bölsterli, René Hvidberg Petersen and their teams are thanked for plant care. Erwin Schoof and the proteomics platform at Denmark’s Technical University are thanked for running protein identification by liquid chromatography-mass spectrometry. This work was supported by a Hallas-Møller Ascending Investigator Fellowship grant from Novo Nordisk Foundation (NNF19OC0054973), a Consolidator Grant from the European Research Council (PATHORISC, ERC-2016-CoG 726417), a Research Infrastructure Grant from Carlsberg Fondet (CF20-0659), and an Instrument Grant from Brdr Hartmann Fonden (A35879), all to P.B.

## AUTHOR CONTRIBUTIONS

**Mathias Due Tankmar:** Investigation; formal analysis; visualization; writing – review and editing. **Marlene Reichel:** Investigation; writing – review and editing. **Laura Arribas-Hernández:** Conceptualization; supervision; visualization; writing – review and editing. **Peter Brodersen:** Conceptualization; supervision; funding acquisition; writing – original draft.

---

**Figure EV1.**
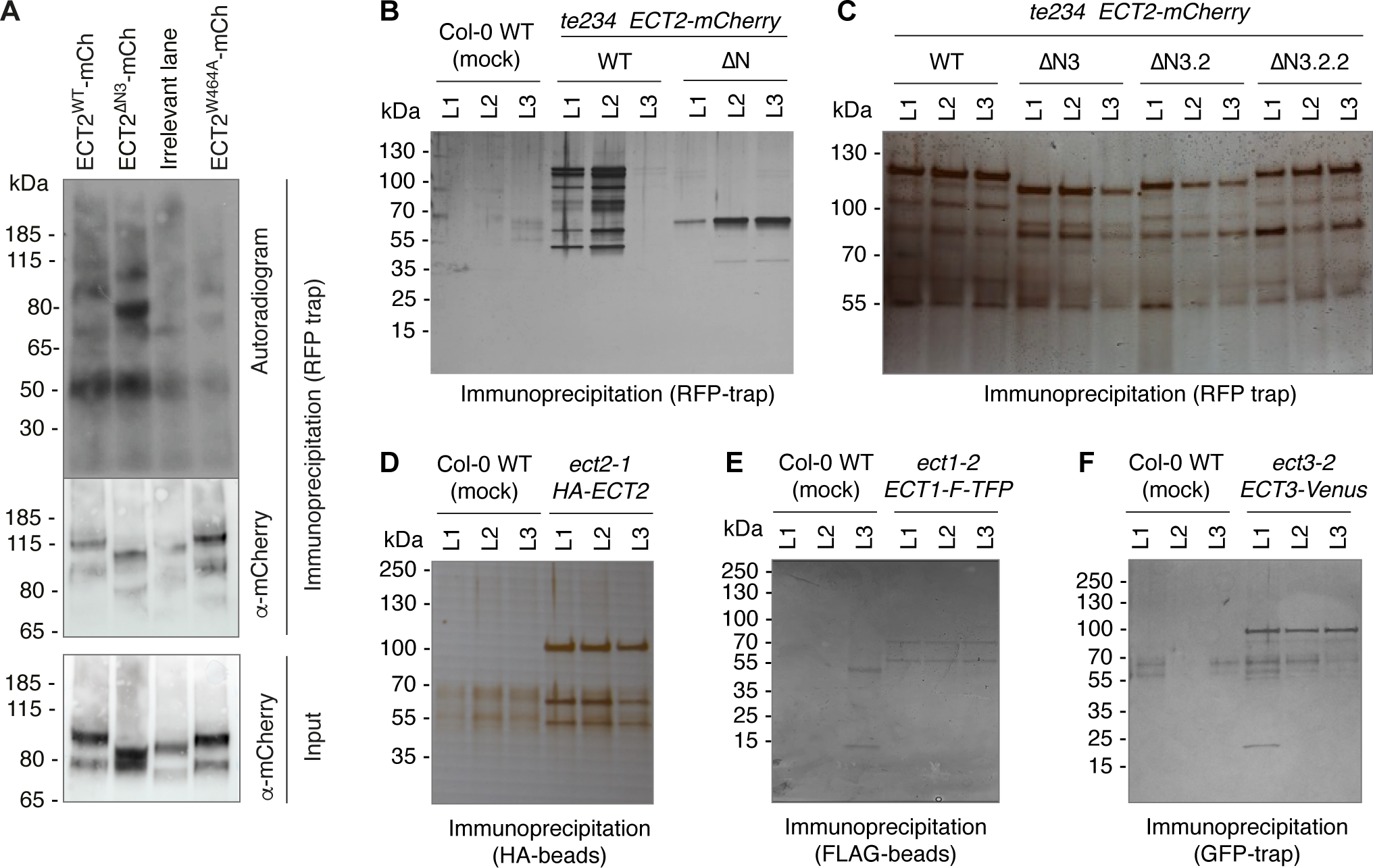
The N3 element is necessary for interaction between ECT2 and PAB2/4/8 (extended data) A Uncropped gel of the CLIP analysis in Fig 3A. B-F Silver-stained gels showing the immunoprecipitations for the MS-IP experiments in Fig 3B-D and F-H, and 4E.

**Figure EV2.**
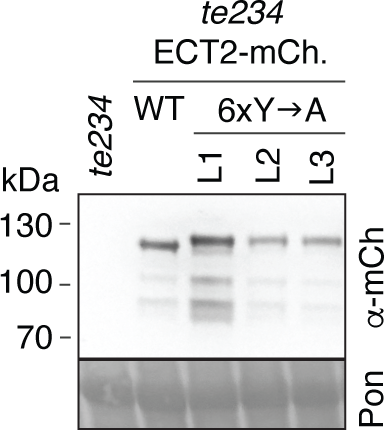
Expression of ECT2^6xY➔A^-mCherry. Western blot showing the amount of ECT2^6xY➔A^-mCherry protein in T2 seedlings of three independent transgenic lines (L1-L3) compared to wild type ECT2-mCherry. Ponceau (Pon) staining is used as loading control.

**Figure EV3.**
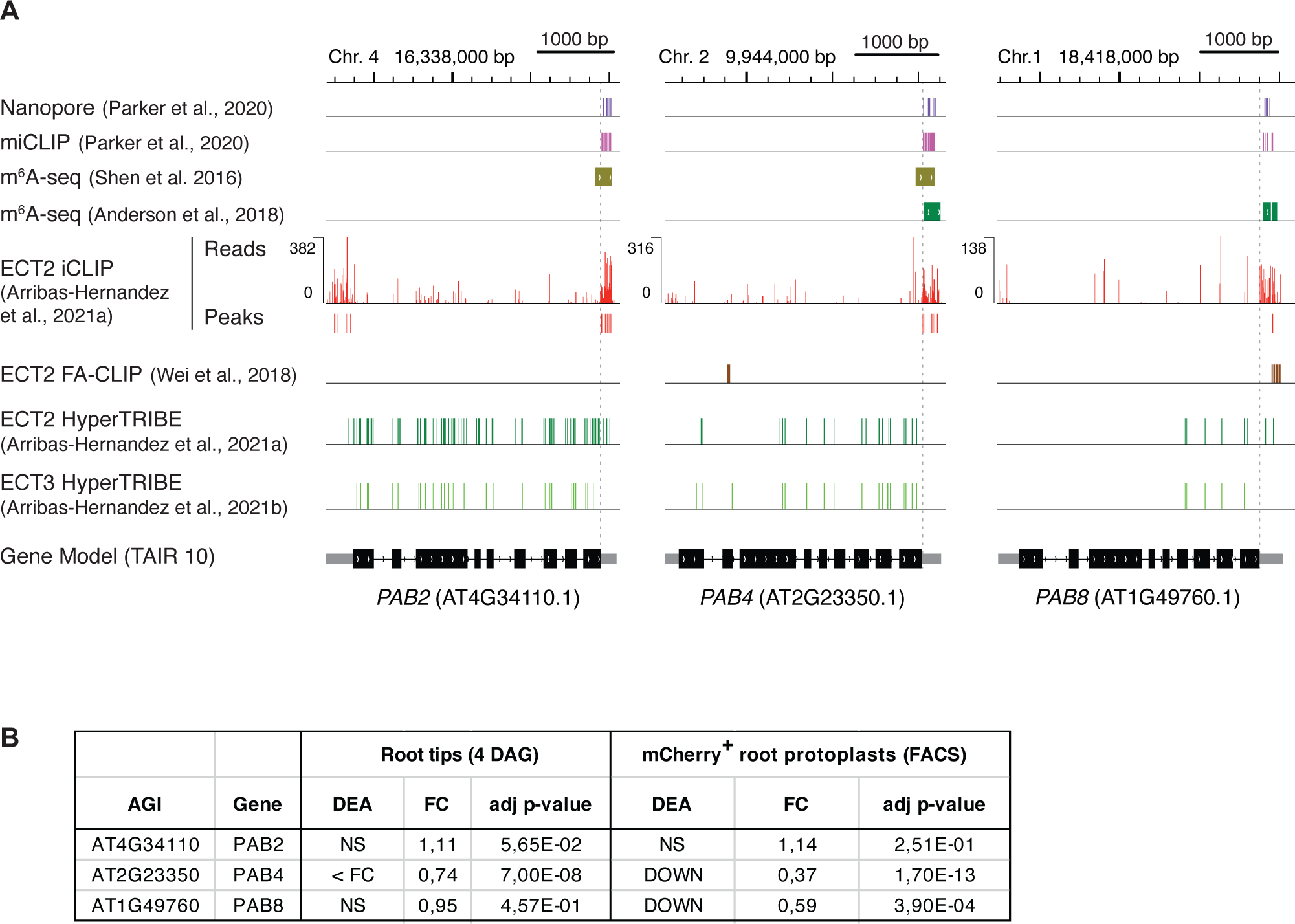
PAB2/4/8 mRNAs contain m^6^A and are ECT2/3 targets. A Integrative Genomics Viewer (IGV)-representation of *PAB2/4/8* loci showing the position of m^6^A marks mapped by different methods in seedlings (Nanopore (Parker et al., 2020), miCLIP (Parker et al., 2020) and m^6^A-seq (Shen et al, 2016)) and rosettes (m^6^A-seq (Anderson et al, 2018)), ECT2-binding sites determined by iCLIP (Arribas-Hernández et al., 2021a) and FA-CLIP (Wei et al., 2018), and ECT2/ECT3-ADAR-editing sites (HyperTRIBE) (Arribas-Hernández et al., 2021a, Arribas-Hernández et al., 2021b) in seedlings. In the gene models (bottom), boxes represent exons and lines are introns. Black-coloured exons correspond to the coding sequence, and grey is used for the untranslated regions (UTRs). Vertical dotted lines mark the start of 3’UTRs. B Differential abundance of *PAB2/4/8* mRNAs in root tips or rapidly dividing cells that normally express *ECT2* (mCherry+ FACS-sorted root protoplasts), between plants with or without ECT2/3/4 function (Arribas-Hernández et al., 2021b).

