## Supplementary material for "A YTHDF-PABP axis is required for m^6^A-mediated organogenesis in plants": Data Set EV1

**Dataset EV1**

Amino acid sequence of Arabidopsis thaliana ECT2 with the deleted (∆X) or mutated (6xY🡪A) amino acids marked in red for each of the indicated mutants. The YTH domain is coloured blue in all sequences as a reference. The mutants are organized according to figures in the manuscript.

**Figure 1**

∆C

MATVAPPADQATDLLQKLSLDSPAKASEIPEPNKKTAVYQYGGVDVHGQVPSYDRSLTPMLPSDAADPSVCYVPNPYNPYQYYNVYGSGQEWTDYPAYTNPEGVDMNSGIYGENGTVVYPQGYGYAAYPYSPATSPAPQLGGEGQLYGAQQYQYPNYFPNSGPYASSVATPTQPDLSANKPAGVKTLPADSNNVASAAGITKGSNGSAPVKPTNQATLNTSSNLYGMGAPGGGLAAGYQDPRYAYEGYYAPVPWHDGSKYSDVQRPVSGSGVASSYSKSSTVPSSRNQNYRSNSHYTSVHQPSSVTGYGTAQGYYNRMYQNKLYGQYGSTGRSALGYGSSGYDSRTNGRGWAATDNKYRSWGRGNSYYYGNENNVDGLNELNRGPRAKGTKNQKGNLDDSLEVKEQTGESNVTEVGEADNTCVVPDREQYNKEDFPVDYANAMFFIIKSYSEDDVHKSIKYNVAASTPNGNKKLAAAYQEAQQKAGGCPIFLFFSVNASGQFVGLAEMTGPVDFNTNVEYWQQDKWTGSFPLKWHIVKDVPNSLLKHITLENNENKPVTNSRDTQEVKLEQGLKIVKIFKEHSSKTCILDDFSFYEVRQKTILEKKAKQTQKQVSEEKVTDEKKESATAESASKESPAAVQTSSDVKVAENGSVAKPVTGDVVANGC

∆N

MATVAPPADQATDLLQKLSLDSPAKASEIPEPNKKTAVYQYGGVDVHGQVPSYDRSLTPMLPSDAADPSVCYVPNPYNPYQYYNVYGSGQEWTDYPAYTNPEGVDMNSGIYGENGTVVYPQGYGYAAYPYSPATSPAPQLGGEGQLYGAQQYQYPNYFPNSGPYASSVATPTQPDLSANKPAGVKTLPADSNNVASAAGITKGSNGSAPVKPTNQATLNTSSNLYGMGAPGGGLAAGYQDPRYAYEGYYAPVPWHDGSKYSDVQRPVSGSGVASSYSKSSTVPSSRNQNYRSNSHYTSVHQPSSVTGYGTAQGYYNRMYQNKLYGQYGSTGRSALGYGSSGYDSRTNGRGWAATDNKYRSWGRGNSYYYGNENNVDGLNELNRGPRAKGTKNQKGNLDDSLEVKEQTGESNVTEVGEADNTCVVPDREQYNKEDFPVDYANAMFFIIKSYSEDDVHKSIKYNVAASTPNGNKKLAAAYQEAQQKAGGCPIFLFFSVNASGQFVGLAEMTGPVDFNTNVEYWQQDKWTGSFPLKWHIVKDVPNSLLKHITLENNENKPVTNSRDTQEVKLEQGLKIVKIFKEHSSKTCILDDFSFYEVRQKTILEKKAKQTQKQVSEEKVTDEKKESATAESASKESPAAVQTSSDVKVAENGSVAKPVTGDVVANGC

∆N.1

MATVAPPADQATDLLQKLSLDSPAKASEIPEPNKKTAVYQYGGVDVHGQVPSYDRSLTPMLPSDAADPSVCYVPNPYNPYQYYNVYGSGQEWTDYPAYTNPEGVDMNSGIYGENGTVVYPQGYGYAAYPYSPATSPAPQLGGEGQLYGAQQYQYPNYFPNSGPYASSVATPTQPDLSANKPAGVKTLPADSNNVASAAGITKGSNGSAPVKPTNQATLNTSSNLYGMGAPGGGLAAGYQDPRYAYEGYYAPVPWHDGSKYSDVQRPVSGSGVASSYSKSSTVPSSRNQNYRSNSHYTSVHQPSSVTGYGTAQGYYNRMYQNKLYGQYGSTGRSALGYGSSGYDSRTNGRGWAATDNKYRSWGRGNSYYYGNENNVDGLNELNRGPRAKGTKNQKGNLDDSLEVKEQTGESNVTEVGEADNTCVVPDREQYNKEDFPVDYANAMFFIIKSYSEDDVHKSIKYNVAASTPNGNKKLAAAYQEAQQKAGGCPIFLFFSVNASGQFVGLAEMTGPVDFNTNVEYWQQDKWTGSFPLKWHIVKDVPNSLLKHITLENNENKPVTNSRDTQEVKLEQGLKIVKIFKEHSSKTCILDDFSFYEVRQKTILEKKAKQTQKQVSEEKVTDEKKESATAESASKESPAAVQTSSDVKVAENGSVAKPVTGDVVANGC

∆N.2

MATVAPPADQATDLLQKLSLDSPAKASEIPEPNKKTAVYQYGGVDVHGQVPSYDRSLTPMLPSDAADPSVCYVPNPYNPYQYYNVYGSGQEWTDYPAYTNPEGVDMNSGIYGENGTVVYPQGYGYAAYPYSPATSPAPQLGGEGQLYGAQQYQYPNYFPNSGPYASSVATPTQPDLSANKPAGVKTLPADSNNVASAAGITKGSNGSAPVKPTNQATLNTSSNLYGMGAPGGGLAAGYQDPRYAYEGYYAPVPWHDGSKYSDVQRPVSGSGVASSYSKSSTVPSSRNQNYRSNSHYTSVHQPSSVTGYGTAQGYYNRMYQNKLYGQYGSTGRSALGYGSSGYDSRTNGRGWAATDNKYRSWGRGNSYYYGNENNVDGLNELNRGPRAKGTKNQKGNLDDSLEVKEQTGESNVTEVGEADNTCVVPDREQYNKEDFPVDYANAMFFIIKSYSEDDVHKSIKYNVAASTPNGNKKLAAAYQEAQQKAGGCPIFLFFSVNASGQFVGLAEMTGPVDFNTNVEYWQQDKWTGSFPLKWHIVKDVPNSLLKHITLENNENKPVTNSRDTQEVKLEQGLKIVKIFKEHSSKTCILDDFSFYEVRQKTILEKKAKQTQKQVSEEKVTDEKKESATAESASKESPAAVQTSSDVKVAENGSVAKPVTGDVVANGC

**Figure 2**

∆N3

MATVAPPADQATDLLQKLSLDSPAKASEIPEPNKKTAVYQYGGVDVHGQVPSYDRSLTPMLPSDAADPSVCYVPNPYNPYQYYNVYGSGQEWTDYPAYTNPEGVDMNSGIYGENGTVVYPQGYGYAAYPYSPATSPAPQLGGEGQLYGAQQYQYPNYFPNSGPYASSVATPTQPDLSANKPAGVKTLPADSNNVASAAGITKGSNGSAPVKPTNQATLNTSSNLYGMGAPGGGLAAGYQDPRYAYEGYYAPVPWHDGSKYSDVQRPVSGSGVASSYSKSSTVPSSRNQNYRSNSHYTSVHQPSSVTGYGTAQGYYNRMYQNKLYGQYGSTGRSALGYGSSGYDSRTNGRGWAATDNKYRSWGRGNSYYYGNENNVDGLNELNRGPRAKGTKNQKGNLDDSLEVKEQTGESNVTEVGEADNTCVVPDREQYNKEDFPVDYANAMFFIIKSYSEDDVHKSIKYNVAASTPNGNKKLAAAYQEAQQKAGGCPIFLFFSVNASGQFVGLAEMTGPVDFNTNVEYWQQDKWTGSFPLKWHIVKDVPNSLLKHITLENNENKPVTNSRDTQEVKLEQGLKIVKIFKEHSSKTCILDDFSFYEVRQKTILEKKAKQTQKQVSEEKVTDEKKESATAESASKESPAAVQTSSDVKVAENGSVAKPVTGDVVANGC

∆N4

MATVAPPADQATDLLQKLSLDSPAKASEIPEPNKKTAVYQYGGVDVHGQVPSYDRSLTPMLPSDAADPSVCYVPNPYNPYQYYNVYGSGQEWTDYPAYTNPEGVDMNSGIYGENGTVVYPQGYGYAAYPYSPATSPAPQLGGEGQLYGAQQYQYPNYFPNSGPYASSVATPTQPDLSANKPAGVKTLPADSNNVASAAGITKGSNGSAPVKPTNQATLNTSSNLYGMGAPGGGLAAGYQDPRYAYEGYYAPVPWHDGSKYSDVQRPVSGSGVASSYSKSSTVPSSRNQNYRSNSHYTSVHQPSSVTGYGTAQGYYNRMYQNKLYGQYGSTGRSALGYGSSGYDSRTNGRGWAATDNKYRSWGRGNSYYYGNENNVDGLNELNRGPRAKGTKNQKGNLDDSLEVKEQTGESNVTEVGEADNTCVVPDREQYNKEDFPVDYANAMFFIIKSYSEDDVHKSIKYNVAASTPNGNKKLAAAYQEAQQKAGGCPIFLFFSVNASGQFVGLAEMTGPVDFNTNVEYWQQDKWTGSFPLKWHIVKDVPNSLLKHITLENNENKPVTNSRDTQEVKLEQGLKIVKIFKEHSSKTCILDDFSFYEVRQKTILEKKAKQTQKQVSEEKVTDEKKESATAESASKESPAAVQTSSDVKVAENGSVAKPVTGDVVANGC

∆N5

MATVAPPADQATDLLQKLSLDSPAKASEIPEPNKKTAVYQYGGVDVHGQVPSYDRSLTPMLPSDAADPSVCYVPNPYNPYQYYNVYGSGQEWTDYPAYTNPEGVDMNSGIYGENGTVVYPQGYGYAAYPYSPATSPAPQLGGEGQLYGAQQYQYPNYFPNSGPYASSVATPTQPDLSANKPAGVKTLPADSNNVASAAGITKGSNGSAPVKPTNQATLNTSSNLYGMGAPGGGLAAGYQDPRYAYEGYYAPVPWHDGSKYSDVQRPVSGSGVASSYSKSSTVPSSRNQNYRSNSHYTSVHQPSSVTGYGTAQGYYNRMYQNKLYGQYGSTGRSALGYGSSGYDSRTNGRGWAATDNKYRSWGRGNSYYYGNENNVDGLNELNRGPRAKGTKNQKGNLDDSLEVKEQTGESNVTEVGEADNTCVVPDREQYNKEDFPVDYANAMFFIIKSYSEDDVHKSIKYNVAASTPNGNKKLAAAYQEAQQKAGGCPIFLFFSVNASGQFVGLAEMTGPVDFNTNVEYWQQDKWTGSFPLKWHIVKDVPNSLLKHITLENNENKPVTNSRDTQEVKLEQGLKIVKIFKEHSSKTCILDDFSFYEVRQKTILEKKAKQTQKQVSEEKVTDEKKESATAESASKESPAAVQTSSDVKVAENGSVAKPVTGDVVANGC

∆N6

MATVAPPADQATDLLQKLSLDSPAKASEIPEPNKKTAVYQYGGVDVHGQVPSYDRSLTPMLPSDAADPSVCYVPNPYNPYQYYNVYGSGQEWTDYPAYTNPEGVDMNSGIYGENGTVVYPQGYGYAAYPYSPATSPAPQLGGEGQLYGAQQYQYPNYFPNSGPYASSVATPTQPDLSANKPAGVKTLPADSNNVASAAGITKGSNGSAPVKPTNQATLNTSSNLYGMGAPGGGLAAGYQDPRYAYEGYYAPVPWHDGSKYSDVQRPVSGSGVASSYSKSSTVPSSRNQNYRSNSHYTSVHQPSSVTGYGTAQGYYNRMYQNKLYGQYGSTGRSALGYGSSGYDSRTNGRGWAATDNKYRSWGRGNSYYYGNENNVDGLNELNRGPRAKGTKNQKGNLDDSLEVKEQTGESNVTEVGEADNTCVVPDREQYNKEDFPVDYANAMFFIIKSYSEDDVHKSIKYNVAASTPNGNKKLAAAYQEAQQKAGGCPIFLFFSVNASGQFVGLAEMTGPVDFNTNVEYWQQDKWTGSFPLKWHIVKDVPNSLLKHITLENNENKPVTNSRDTQEVKLEQGLKIVKIFKEHSSKTCILDDFSFYEVRQKTILEKKAKQTQKQVSEEKVTDEKKESATAESASKESPAAVQTSSDVKVAENGSVAKPVTGDVVANGC

∆N7

MATVAPPADQATDLLQKLSLDSPAKASEIPEPNKKTAVYQYGGVDVHGQVPSYDRSLTPMLPSDAADPSVCYVPNPYNPYQYYNVYGSGQEWTDYPAYTNPEGVDMNSGIYGENGTVVYPQGYGYAAYPYSPATSPAPQLGGEGQLYGAQQYQYPNYFPNSGPYASSVATPTQPDLSANKPAGVKTLPADSNNVASAAGITKGSNGSAPVKPTNQATLNTSSNLYGMGAPGGGLAAGYQDPRYAYEGYYAPVPWHDGSKYSDVQRPVSGSGVASSYSKSSTVPSSRNQNYRSNSHYTSVHQPSSVTGYGTAQGYYNRMYQNKLYGQYGSTGRSALGYGSSGYDSRTNGRGWAATDNKYRSWGRGNSYYYGNENNVDGLNELNRGPRAKGTKNQKGNLDDSLEVKEQTGESNVTEVGEADNTCVVPDREQYNKEDFPVDYANAMFFIIKSYSEDDVHKSIKYNVAASTPNGNKKLAAAYQEAQQKAGGCPIFLFFSVNASGQFVGLAEMTGPVDFNTNVEYWQQDKWTGSFPLKWHIVKDVPNSLLKHITLENNENKPVTNSRDTQEVKLEQGLKIVKIFKEHSSKTCILDDFSFYEVRQKTILEKKAKQTQKQVSEEKVTDEKKESATAESASKESPAAVQTSSDVKVAENGSVAKPVTGDVVANGC

∆N8

MATVAPPADQATDLLQKLSLDSPAKASEIPEPNKKTAVYQYGGVDVHGQVPSYDRSLTPMLPSDAADPSVCYVPNPYNPYQYYNVYGSGQEWTDYPAYTNPEGVDMNSGIYGENGTVVYPQGYGYAAYPYSPATSPAPQLGGEGQLYGAQQYQYPNYFPNSGPYASSVATPTQPDLSANKPAGVKTLPADSNNVASAAGITKGSNGSAPVKPTNQATLNTSSNLYGMGAPGGGLAAGYQDPRYAYEGYYAPVPWHDGSKYSDVQRPVSGSGVASSYSKSSTVPSSRNQNYRSNSHYTSVHQPSSVTGYGTAQGYYNRMYQNKLYGQYGSTGRSALGYGSSGYDSRTNGRGWAATDNKYRSWGRGNSYYYGNENNVDGLNELNRGPRAKGTKNQKGNLDDSLEVKEQTGESNVTEVGEADNTCVVPDREQYNKEDFPVDYANAMFFIIKSYSEDDVHKSIKYNVAASTPNGNKKLAAAYQEAQQKAGGCPIFLFFSVNASGQFVGLAEMTGPVDFNTNVEYWQQDKWTGSFPLKWHIVKDVPNSLLKHITLENNENKPVTNSRDTQEVKLEQGLKIVKIFKEHSSKTCILDDFSFYEVRQKTILEKKAKQTQKQVSEEKVTDEKKESATAESASKESPAAVQTSSDVKVAENGSVAKPVTGDVVANGC

∆N3.1

MATVAPPADQATDLLQKLSLDSPAKASEIPEPNKKTAVYQYGGVDVHGQVPSYDRSLTPMLPSDAADPSVCYVPNPYNPYQYYNVYGSGQEWTDYPAYTNPEGVDMNSGIYGENGTVVYPQGYGYAAYPYSPATSPAPQLGGEGQLYGAQQYQYPNYFPNSGPYASSVATPTQPDLSANKPAGVKTLPADSNNVASAAGITKGSNGSAPVKPTNQATLNTSSNLYGMGAPGGGLAAGYQDPRYAYEGYYAPVPWHDGSKYSDVQRPVSGSGVASSYSKSSTVPSSRNQNYRSNSHYTSVHQPSSVTGYGTAQGYYNRMYQNKLYGQYGSTGRSALGYGSSGYDSRTNGRGWAATDNKYRSWGRGNSYYYGNENNVDGLNELNRGPRAKGTKNQKGNLDDSLEVKEQTGESNVTEVGEADNTCVVPDREQYNKEDFPVDYANAMFFIIKSYSEDDVHKSIKYNVAASTPNGNKKLAAAYQEAQQKAGGCPIFLFFSVNASGQFVGLAEMTGPVDFNTNVEYWQQDKWTGSFPLKWHIVKDVPNSLLKHITLENNENKPVTNSRDTQEVKLEQGLKIVKIFKEHSSKTCILDDFSFYEVRQKTILEKKAKQTQKQVSEEKVTDEKKESATAESASKESPAAVQTSSDVKVAENGSVAKPVTGDVVANGC

∆N3.2

MATVAPPADQATDLLQKLSLDSPAKASEIPEPNKKTAVYQYGGVDVHGQVPSYDRSLTPMLPSDAADPSVCYVPNPYNPYQYYNVYGSGQEWTDYPAYTNPEGVDMNSGIYGENGTVVYPQGYGYAAYPYSPATSPAPQLGGEGQLYGAQQYQYPNYFPNSGPYASSVATPTQPDLSANKPAGVKTLPADSNNVASAAGITKGSNGSAPVKPTNQATLNTSSNLYGMGAPGGGLAAGYQDPRYAYEGYYAPVPWHDGSKYSDVQRPVSGSGVASSYSKSSTVPSSRNQNYRSNSHYTSVHQPSSVTGYGTAQGYYNRMYQNKLYGQYGSTGRSALGYGSSGYDSRTNGRGWAATDNKYRSWGRGNSYYYGNENNVDGLNELNRGPRAKGTKNQKGNLDDSLEVKEQTGESNVTEVGEADNTCVVPDREQYNKEDFPVDYANAMFFIIKSYSEDDVHKSIKYNVAASTPNGNKKLAAAYQEAQQKAGGCPIFLFFSVNASGQFVGLAEMTGPVDFNTNVEYWQQDKWTGSFPLKWHIVKDVPNSLLKHITLENNENKPVTNSRDTQEVKLEQGLKIVKIFKEHSSKTCILDDFSFYEVRQKTILEKKAKQTQKQVSEEKVTDEKKESATAESASKESPAAVQTSSDVKVAENGSVAKPVTGDVVANGC

∆N8/3.2

MATVAPPADQATDLLQKLSLDSPAKASEIPEPNKKTAVYQYGGVDVHGQVPSYDRSLTPMLPSDAADPSVCYVPNPYNPYQYYNVYGSGQEWTDYPAYTNPEGVDMNSGIYGENGTVVYPQGYGYAAYPYSPATSPAPQLGGEGQLYGAQQYQYPNYFPNSGPYASSVATPTQPDLSANKPAGVKTLPADSNNVASAAGITKGSNGSAPVKPTNQATLNTSSNLYGMGAPGGGLAAGYQDPRYAYEGYYAPVPWHDGSKYSDVQRPVSGSGVASSYSKSSTVPSSRNQNYRSNSHYTSVHQPSSVTGYGTAQGYYNRMYQNKLYGQYGSTGRSALGYGSSGYDSRTNGRGWAATDNKYRSWGRGNSYYYGNENNVDGLNELNRGPRAKGTKNQKGNLDDSLEVKEQTGESNVTEVGEADNTCVVPDREQYNKEDFPVDYANAMFFIIKSYSEDDVHKSIKYNVAASTPNGNKKLAAAYQEAQQKAGGCPIFLFFSVNASGQFVGLAEMTGPVDFNTNVEYWQQDKWTGSFPLKWHIVKDVPNSLLKHITLENNENKPVTNSRDTQEVKLEQGLKIVKIFKEHSSKTCILDDFSFYEVRQKTILEKKAKQTQKQVSEEKVTDEKKESATAESASKESPAAVQTSSDVKVAENGSVAKPVTGDVVANGC

**Figure 4**

∆N3.2.1

MATVAPPADQATDLLQKLSLDSPAKASEIPEPNKKTAVYQYGGVDVHGQVPSYDRSLTPMLPSDAADPSVCYVPNPYNPYQYYNVYGSGQEWTDYPAYTNPEGVDMNSGIYGENGTVVYPQGYGYAAYPYSPATSPAPQLGGEGQLYGAQQYQYPNYFPNSGPYASSVATPTQPDLSANKPAGVKTLPADSNNVASAAGITKGSNGSAPVKPTNQATLNTSSNLYGMGAPGGGLAAGYQDPRYAYEGYYAPVPWHDGSKYSDVQRPVSGSGVASSYSKSSTVPSSRNQNYRSNSHYTSVHQPSSVTGYGTAQGYYNRMYQNKLYGQYGSTGRSALGYGSSGYDSRTNGRGWAATDNKYRSWGRGNSYYYGNENNVDGLNELNRGPRAKGTKNQKGNLDDSLEVKEQTGESNVTEVGEADNTCVVPDREQYNKEDFPVDYANAMFFIIKSYSEDDVHKSIKYNVAASTPNGNKKLAAAYQEAQQKAGGCPIFLFFSVNASGQFVGLAEMTGPVDFNTNVEYWQQDKWTGSFPLKWHIVKDVPNSLLKHITLENNENKPVTNSRDTQEVKLEQGLKIVKIFKEHSSKTCILDDFSFYEVRQKTILEKKAKQTQKQVSEEKVTDEKKESATAESASKESPAAVQTSSDVKVAENGSVAKPVTGDVVANGC

∆N3.2.2

MATVAPPADQATDLLQKLSLDSPAKASEIPEPNKKTAVYQYGGVDVHGQVPSYDRSLTPMLPSDAADPSVCYVPNPYNPYQYYNVYGSGQEWTDYPAYTNPEGVDMNSGIYGENGTVVYPQGYGYAAYPYSPATSPAPQLGGEGQLYGAQQYQYPNYFPNSGPYASSVATPTQPDLSANKPAGVKTLPADSNNVASAAGITKGSNGSAPVKPTNQATLNTSSNLYGMGAPGGGLAAGYQDPRYAYEGYYAPVPWHDGSKYSDVQRPVSGSGVASSYSKSSTVPSSRNQNYRSNSHYTSVHQPSSVTGYGTAQGYYNRMYQNKLYGQYGSTGRSALGYGSSGYDSRTNGRGWAATDNKYRSWGRGNSYYYGNENNVDGLNELNRGPRAKGTKNQKGNLDDSLEVKEQTGESNVTEVGEADNTCVVPDREQYNKEDFPVDYANAMFFIIKSYSEDDVHKSIKYNVAASTPNGNKKLAAAYQEAQQKAGGCPIFLFFSVNASGQFVGLAEMTGPVDFNTNVEYWQQDKWTGSFPLKWHIVKDVPNSLLKHITLENNENKPVTNSRDTQEVKLEQGLKIVKIFKEHSSKTCILDDFSFYEVRQKTILEKKAKQTQKQVSEEKVTDEKKESATAESASKESPAAVQTSSDVKVAENGSVAKPVTGDVVANGC

6xY🡪A MATVAPPADQATDLLQKLSLDSPAKASEIPEPNKKTAVYQYGGVDVHGQVPSYDRSLTPMLPSDAADPSVCYVPNPYNPYQYYNVYGSGQEWTDYPAYTNPEGVDMNSGIYGENGTVVYPQGYGYAAYPYSPATSPAPQLGGEGQLYGAQQYQYPNYFPNSGPYASSVATPTQPDLSANKPAGVKTLPADSNNVASAAGITKGSNGSAPVKPTNQATLNTSSNLYGMGAPGGGLAAGYQDPRYAYEGYYAPVPWHDGSKYSDVQRPVSGSGVASSYSKSSTVPSSRNQNYRSNSHYTSVHQPSSVTGYGTAQGYYNRMYQNKLYGQYGSTGRSALGYGSSGYDSRTNGRGWAATDNKYRSWGRGNSYYYGNENNVDGLNELNRGPRAKGTKNQKGNLDDSLEVKEQTGESNVTEVGEADNTCVVPDREQYNKEDFPVDYANAMFFIIKSYSEDDVHKSIKYNVAASTPNGNKKLAAAYQEAQQKAGGCPIFLFFSVNASGQFVGLAEMTGPVDFNTNVEYWQQDKWTGSFPLKWHIVKDVPNSLLKHITLENNENKPVTNSRDTQEVKLEQGLKIVKIFKEHSSKTCILDDFSFYEVRQKTILEKKAKQTQKQVSEEKVTDEKKESATAESASKESPAAVQTSSDVKVAENGSVAKPVTGDVVANGC
